# Cell Penetrating Thyclotides Facilitate Efficient Delivery of Bioactive Peptides into Cells

**DOI:** 10.64898/2026.07.01.735572

**Authors:** Gamze Ayaz, Hongchao Zheng, Harsha Amarasekara, Victor Clausse, Andy Tran, Ferenc Livák, Michael Kruhlak, Daniel H. Appella

**Author notes:** Correspondence to (D.H.A.). These authors contributed equally to this work (Gamze Ayaz, Hongchao Zheng). Evotec (France) SAS, 195 Route d(Espagne, 31036 Toulouse CEDEX, France.

## Abstract

Cell penetrating thyclotides (CPTs) are synthetic molecules that promote highly efficient cellular uptake and endosomal escape of bioactive peptides. While peptides are valuable as medicinal agents, their translation to therapies is often limited by their inability to cross cell membranes. CPTs have a unique combination of chiral tetrahydrofurans and polar sidechains within a molecular scaffold that can be optimized to efficiently deliver peptide cargo into cells. The cellular uptake and endosomal escape of two peptides with anticancer biological activities but low bioavailabilities were remarkably improved after conjugation to a CPT. Using CPTs to overcome barriers to cellular uptake represents a new direction for the intracellular delivery of bioactive molecules, and will accelerate drug development for new medical therapies.

## Main Text

Intracellular delivery of bioactive molecules is a difficult barrier for many drugs with therapeutic potential.^1,2^ While drugs with molecular weights below 500 Daltons may pass through cell membranes by passive diffusion, larger and more complex molecules often rely on delivery vehicles to access the interior of a cell.^3^ A wide range of delivery vehicles have been investigated, such as polymers, nanoparticles, liposomes, viruses, and cell-penetrating peptides.^4^ These approaches involve combining a biologically active molecule (the cargo) with a delivery vehicle that in many cases results in the vehicle plus cargo being actively transported into an endosome.^5^ Even though the endosome is inside the cell, the molecular cargo must subsequently escape from the endosome and enter the cytosol to exert biological activity.^6^ A common shortcoming of delivery vehicles is that most of the molecular cargo remains trapped inside endosomes where it remains inactive and may ultimately be degraded.^7^ This shortcoming is particularly problematic when the molecular cargo is an α-amino acid polypeptide (a peptide). Techniques to design and identify peptides that potently bind to and inhibit a biological target *in vitro* are highly successful and continue to rapidly improve with the use of artificial intelligence.^8^ Unfortunately, most peptides are unable to enter cells.^9^ To improve the bioavailability of peptides, common strategies include chemical modification with unique amino acids,^10,11^ cyclization via stapled peptides^12^ or related strategies,^13–19^ or the attachment of cell penetrating peptides.^20,21^ These strategies frequently rely on the presence of multiple positive charges from primary amines or guanidines, and cell penetrating peptides commonly have multiple lysines and arginines in their sequences. While cell penetrating peptides can promote cell uptake of peptide cargo, these types of peptides are plagued by problems of stability, non-specific toxicity to healthy cells, and lack of endosomal escape that limit their use for drug development.^22,23^ A completely synthetic molecular scaffold that replaces cell penetrating peptides could overcome these problems and significantly enable the *in vivo* translation of bioactive peptides. However, the synthetic versions of cell penetrating peptides continue to rely on multiple guanidine groups for cell uptake which lead to non-specific toxicity and endosomal entrapment.^24–27^ Identifying new chemical groups that lead to cellular uptake remain very challenging, and there is currently no synthetic platform that can be used as replacement for cell penetrating peptides.

### CPTs enhance the uptake of target peptides for cancer cells

We report a synthetic molecular scaffold called cell penetrating thyclotide (CPT) that effectively delivers peptides into cells (Fig. 1A). CPTs are constructed from a series of amide-linked monomers containing chiral tetrahydrofuran (THF) groups and non-natural polar sidechains (Fig.1B). The sidechains can be easily varied to optimize for cell uptake. Using solid phase peptide synthesis, the CPT monomer units are linked together and easily conjugated to bioactive peptides. Two previously reported peptide inhibitors (peptides **1** and **2**, along with their fluorescein-labeled derivatives Fl-**1** and Fl-**2**, Fig. 1C) were selected for detailed investigations into CPT-promoted cell uptake. In both cases, the peptide inhibitors had been optimized to bind their targets *in vitro* but lacked bioavailability in cells.^28,29^ CPT conjugates of the peptides yielded CPT-**1** and CPT-**2**, and the fluorescein-labeled versions of the conjugates (Fl-CPT-**1** and Fl-CPT-**2**) were also prepared. We initially attached a range of fluorescein-labeled CPTs with different sidechains to **1**, which is a polar peptide that modestly inhibits the enzyme SetD8 (a histone methyl-transferase that is overexpressed in several cancers).^30–32^ From this initial screen (Fig. S1A and B), we identified a piperidine sidechain in the CPT that optimally promotes cellular uptake. When SK-NA-S cells were treated with 5 µM Fl-CPT-**1** for 24 hours, imaging flow cytometry (Amnis ImageStreamX Mark II) (Fig. S1A and B) and confocal microscopy demonstrated remarkably efficient cellular uptake and diffuse distribution throughout the cytosol (Fig. 2A). Approximately 99% of SK-N-AS cells were positive for Fl-CPT-**1**, and among these, 77% exhibited predominantly cytoplasmic localization and 23% showed nuclear distribution (Fig. S2A). In addition, the median fluorescence intensity (MFI) was significantly higher than in cells treated with Fl-**1**. In the absence of CPT, FL-**1** also demonstrated relatively high cellular uptake (85.7% FITC positive cells); however, its MFI was significantly lower than that of Fl-CPT-**1** (Fig. S2, B and C), indicating that only small amounts of FL-**1** enter cells. To determine whether CPT-1 is localized to early endosomes following cellular uptake, SK-N-AS cells were transfected with CellLight™ Early Endosomes-RFP (Rab5a, BacMam 2.0) for 24 h and subsequently incubated with CPT-1 for an additional 1 h. Immunofluorescence analysis showed no detectable co-localization between CPT-1 and the early endosomal marker Rab5a, indicating that CPT-1 is not retained within early endosomes after internalization (Fig. S2D). Although FL-CPT-1 was efficiently internalized without endosomal entrapment, we were unable to observe consistent biological activity for CPT-**1** in cell-based assays possibly due to the modest inhibitory activity or rapid degradation of the peptide once inside the cell. The same CPT was next attached to peptide **2**, a hydrophobic peptide that has been highly optimized as an inhibitor of MDM2 binding to p53.^29^ Overexpression of MDM2 is associated with several cancers and has been a frequent target for therapeutic intervention.^33–35^ The small molecule Nutlin-3 also inhibits MDM2 and is a clinical candidate for the treatment of cancers, but undesired toxic side-effects in clinical trials have limited its development.^36–39^ *In vitro*, peptide **2** binds very strongly to MDM2 and its homolog MDMX, yet it has weak activity in cell-based assays.^29^ We confirmed that fluorescein-labeled peptide **2** (Fl-**2**) does not enter cells (Fig. S2E). In contrast, CPT-**2** conjugates dramatically overcome the inherently poor bioavailability of peptide **2**. As shown in Figure 2B, the fluorescein-labeled version of CPT-**2** (Fl-CPT-**2**) is able penetrate into MCF7 cells with diffuse distribution across the cytosol. Approximately 98% of MCF7 cells internalized the Fl-CPT-**2**, which was predominantly localized in the cytoplasm and partially in the nucleus. In contrast, the Fl-**2** lacking the CPT moiety exhibited minimal cellular uptake, with only approximately 25% of cells showing detectable fluorescence (Fig. S2E). Consistent with these observations, the MFI was significantly higher in cells treated with the Fl-CPT-**2** compared to those treated with the Fl-**2** (Fig. S2, F and G). Similar to CPT-1, CPT-2 also showed no detectable co-localization with the early endosomal marker Rab5a following cellular uptake (fig. S2H). These findings indicate that the CPT moiety may facilitate efficient escape from early endosomes independent of the peptide sequence.

**Figure 1.**
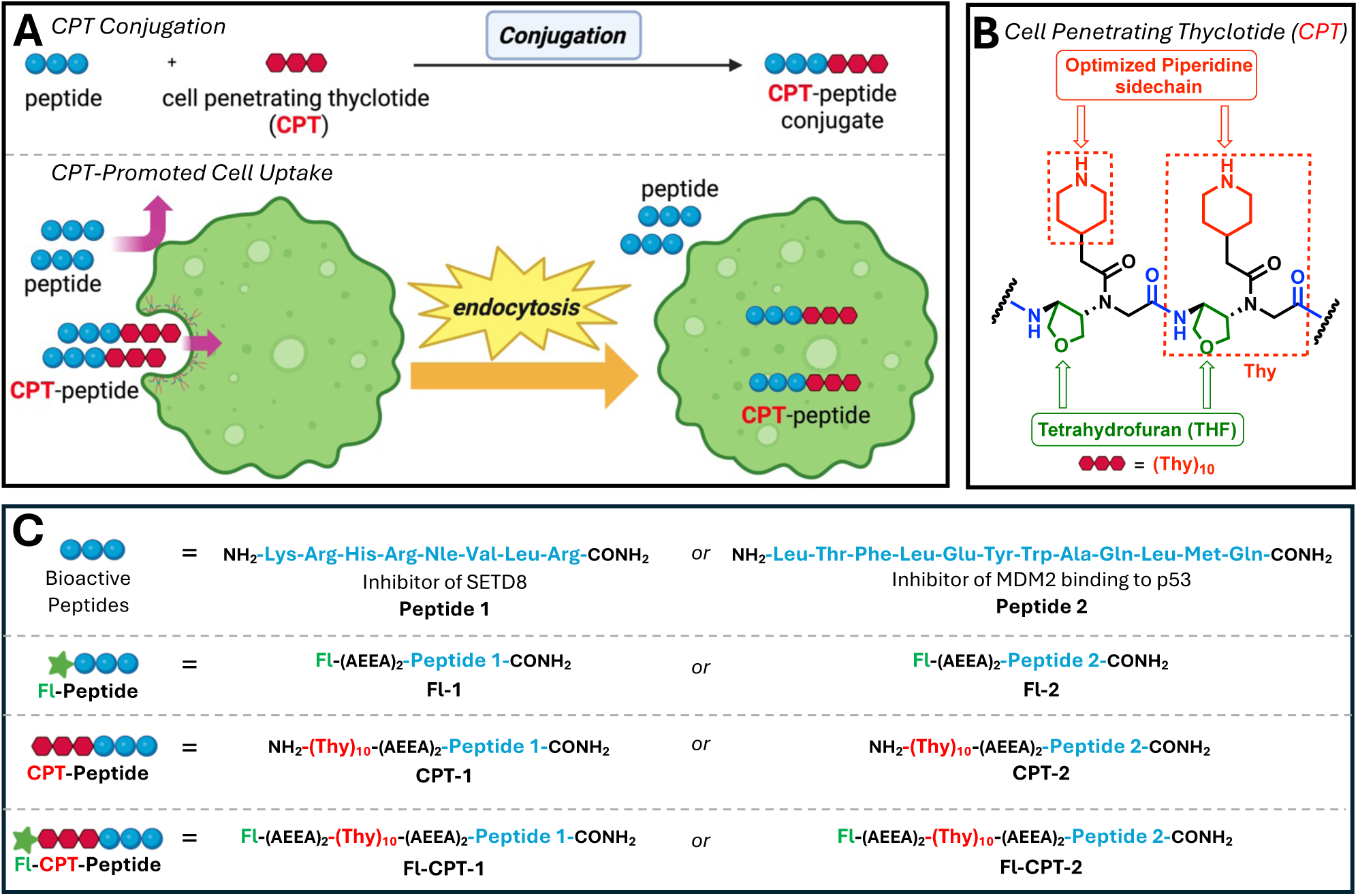
Design and schematic structure of cell penetrating thyclotide (CPT) conjugates. **(A)** Schematic illustration of the proposed cellular uptake mechanism of CPT-peptide conjugates. Following conjugation to cargo peptides, the cell-penetrating thyclotide (CPT) facilitates intracellular delivery into cancer cells primarily through endocytosis-mediated uptake pathways. **(B)** Chemical structure of the CPT monomer used for peptide conjugation. The structural features responsible for cellular internalization and enhanced solubility are highlighted. **(C)** Schematic representation of the peptide conjugates and their corresponding fluorescein-labelled derivatives used in this study. Peptide sequences, CPT-conjugated constructs, and fluorescein (FL) labeled molecules employed for cellular uptake, localization and functional analyses are shown.

**Figure 2.**
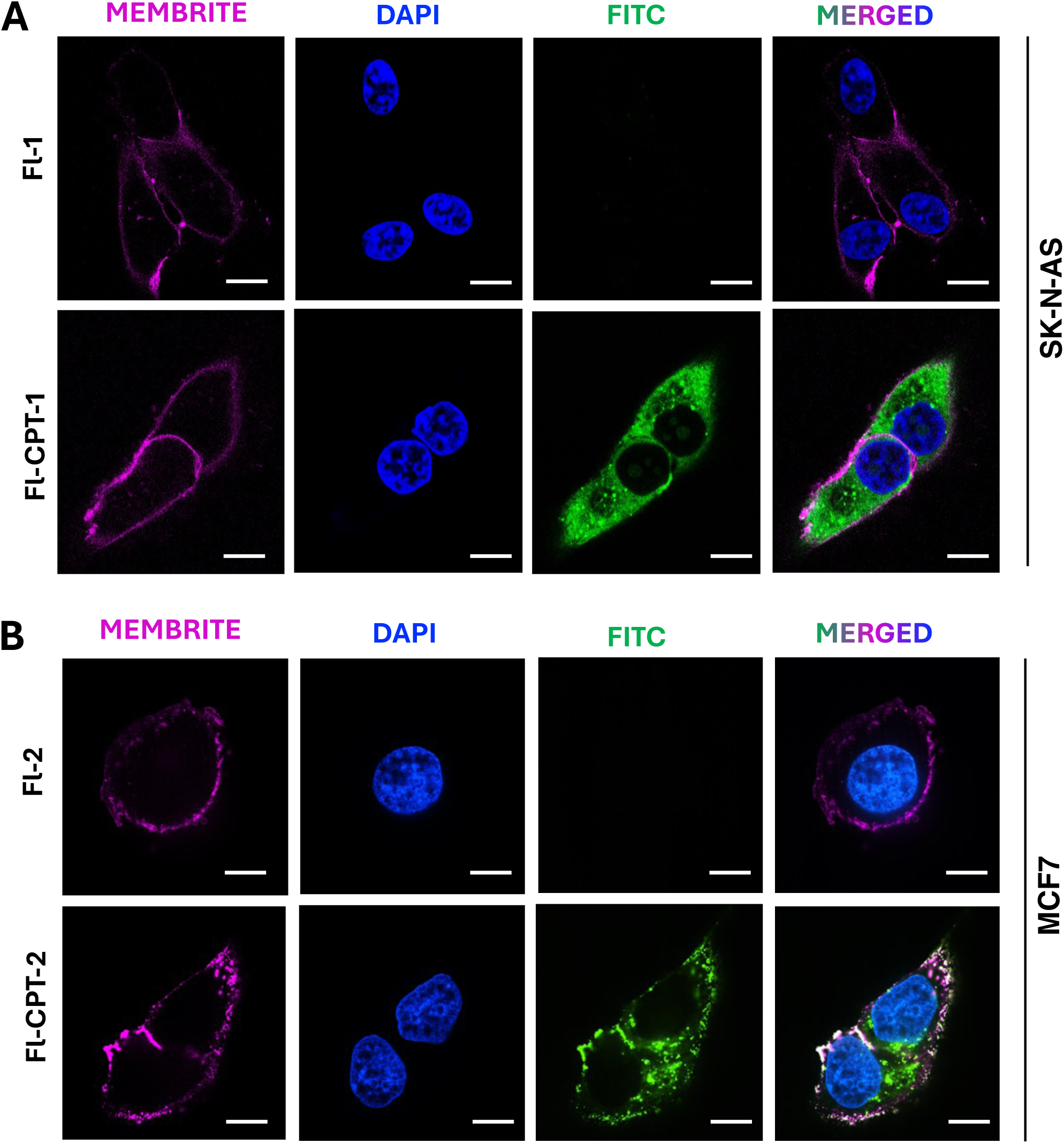
Subcellular localization of CPT conjugated peptides. **(A)** SK-N-AS cells were treated with 5 µM FL-**1** or FL-CPT-**1** for 16h. Cell membrane were stained using the Membrite Fix 640/660 cell surface staining kit, and nuclei were counterstained with DAPI. Scale bar, 10 µm. **(B)** MCF7 cells were treated with 5 µM FL-**2** or FL-CPT-**2** for 16 h. Cell membrane were stained using the Membrite Fix 640/660 cell surface staining kit, and nuclei were counterstained with DAPI. Scale bar, 10 µm.

### Bioavailability of CPT-2 in cancer cells

The biological activity of CPT-**2** was examined in detail to confirm that the molecule inhibits the MDM2-p53 interaction and leads to reactivation of p53. Treatment of MCF7 breast cancer cells, which have wild type p53, with 10 µM of CPT-**2** results in upregulation of p53 and induction of its downstream targets p21 (which leads to cell cycle arrest) and Bax (a proapoptotic protein). The molecule Nutlin-3 was used as a positive control and shows similar effects. In contrast, **2** alone has no effect. Modest increases in MDM2 and MDMX expression were also observed following CPT-**2** treatment (Fig. 3A), also consistent with the activity of Nutlin-3. To demonstrate specificity, control cells (MDA-MB-231) with a R280K mutation in p53 were used. The p53 mutation in the these cells prevents DNA binding and reactivation of p21 and BAX. Therefore, inhibition of MDM2 in MDA-MB-231 cells will not promote increases in p21 or Bax as p53 is unable to function due to the mutation. After confirming that Fl-CPT-**2** was similarly taken up by MDA-MB-231 cells (Fig. S3, A and C), treatment with CPT-**2** and Nutlin-3 both show no effects on the p21 and BAX protein levels (Fig. 3B). To further assess whether stabilized p53 accumulates in the nucleus together with MDM2, we performed a proximity ligation assay (PLA) in treated MCF7 cells. Fl-CPT-2 treatment for 24 h induced prominent nuclear PLA foci, indicating increased p53-MDM2 co-localization compared to the DMSO control (fig. S4, A and B). Because Fl-CPT-**2** does not localize to the nucleus, these foci reflect the accumulation of both interaction partners rather than a direct action of Fl-CPT-**2**. Inhibition of the MDM2/MDMX-p53 interaction stabilizes p53, which translocates to the nucleus and transcriptionally upregulates MDM2 through the p53-MDM2 autoregulatory feedback loop. The increase in proximity signal therefore reflects the combined nuclear accumulation of stabilized p53 and feedback-induced MDM2, which was also reported for Nutlin-3^40,41^.

**Figure 3.**
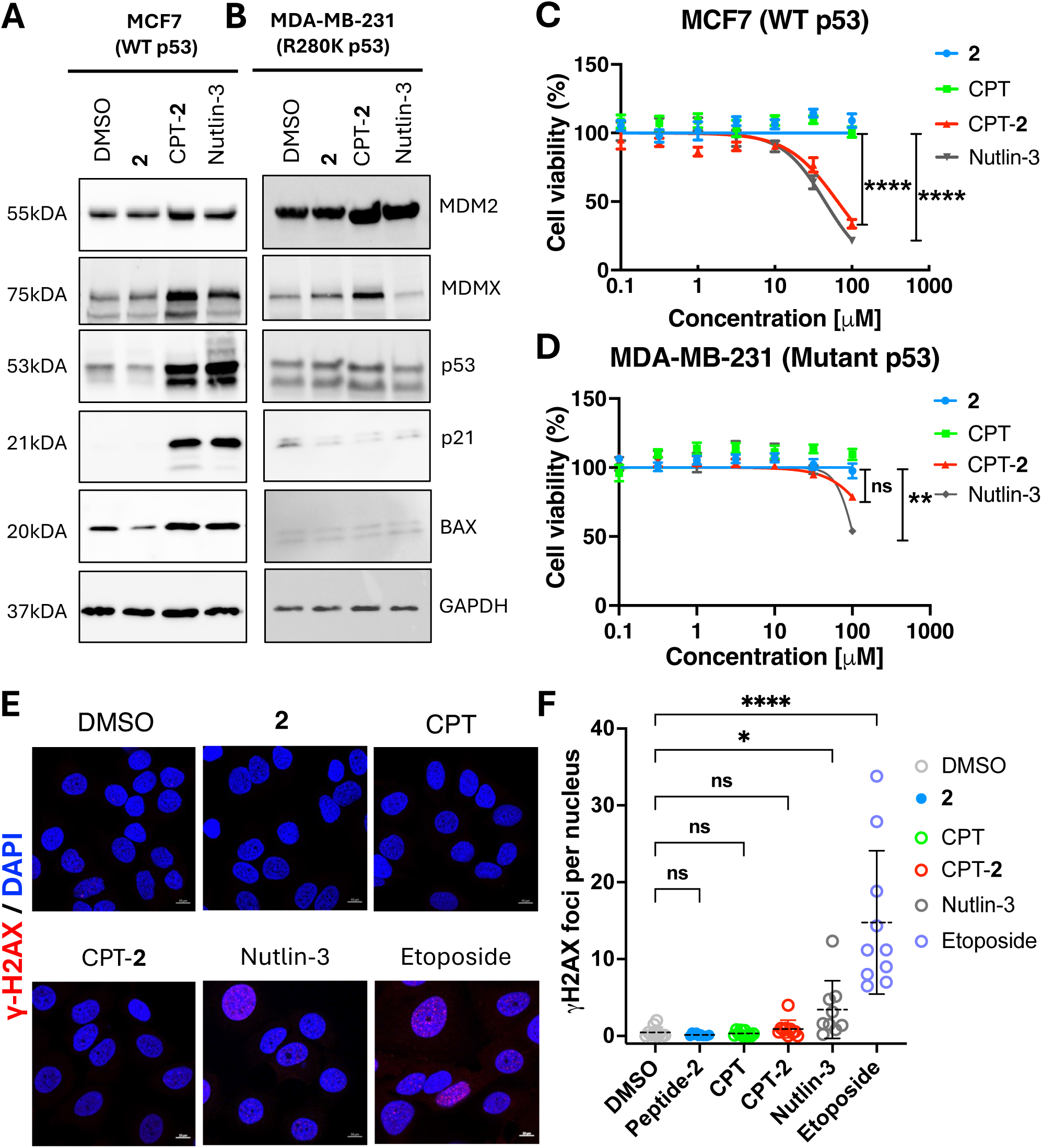
CPT-2 reduces cell viability through wild-type p53-dependent pathways *in vitro.* **(A, B)** Western blot analysis of protein expression in MCF7 **(A)** and MDA-MB-231 **(B)** cells following treatment with the indicated compounds at 10 µM for 24 h. Representative immunoblots showing modulation of p53-associated signaling pathways are presented. **(C, D)** Cell viability of MCF7 **(C)** and MDA-MB-231 **(D)** cells following treatment with the indicated compounds at concentrations ranging from 0.1 to 100 µM for 72 h, as determined by MTS assay. Data are presented as mean ± SEM from three independent biological replicates. Statistical significance was determined using the Kruskal-Wallis test followed by Dunn’s multiple comparisons test. P < 0.05 considered statistically significant. ****P ≤ 0.0001, **P ≤ 0.01, ns; not significant. **(E)** Immunofluorescence analysis of γ-H2AX foci formation in MCF7 cells following treatment with the indicated compounds at 50 µM for 96 h, except for Nutlin-3, which was used at 10 µM. Nuclei were counterstained with DAPI. Increased γ-H2AX staining indicates induction of DNA damage response pathways. Scale bar, 10 µm. **(F)** γ-H2AX foci per nucleus in MCF7 cells across treatment groups. Foci were quantified in Fiji (ImageJ). Each dot represents the mean number of γ-H2AX foci per nucleus for one image (10 images per group; 98 (DMSO), 88 (Peptide-**2**), 107 (CPT), 64 (CPT-2), 68 (Nutlin-3) and 70 (Etoposide) nuclei analyzed per group). Data were analyzed by Kruskal-Wallis test followed by Dunn’s multiple comparisons test versus the DMSO control. ****P ≤ 0.0001, *P= 0.0465, ns; not significant.

Cytotoxicity assays at 72 h demonstrated that CPT-**2** selectively reduced the viability of MCF7 cells, with efficiency similar to Nutlin-3. In contrast, peptide-**2** and CPT alone (**S10**, Supporting Info, Table S3) showed no measurable effects (Fig. 3C). As expected, MDA-MB-231 cells were mostly unaffected by the same series of molecules except for high concentrations of Nutlin-3 that indicated non-specific toxicity (Fig. 3D). To evaluate potential off-target genotoxicity, we quantified nuclear γH2AX foci after 96 h treatment of MCF7 cells with **2**, CPT-**2**, or control molecules at 50 µM. With CPT-**2**, CPT, and **2**, foci counts were indistinguishable from the DMSO vehicle (Fig. 3, E and F). In contrast, the topoisomerase II inhibitor etoposide induced a pronounced increase in γH2AX foci (P < 0.0001), while Nutlin-3 showed a modest but significant increase (P < 0.05). Under the same western blotting conditions, CPT-**2** increased p53, p21 and Bax to levels comparable to Nutlin-3 and etoposide, yet did not elevate γH2AX (Fig. S5). Collectively, these results demonstrate that CPT-**2** activates p53 signaling through MDM2 inhibition in a wild-type p53-dependent manner while exhibiting reduced non-specific toxicity relative to Nutlin-3.

The next series of experiments examined the effect of CPT-**2** on cell cycle distribution and apoptosis to distinguish whether the loss of proliferation caused by CPT-**2** reflects cell cycle arrest or cell death. Active DNA synthesis was assessed by 5-ethynyl-2’-deoxyuridine (EdU) incorporation combined with FxCycle Violet DNA content staining, allowing replicating (EdU^+^, S-phase) cells to be resolved from non-replicating G1 and G2/M populations. Cells treated with DMSO and **2** displayed a prominent EdU^+^ S phase population (36.9% and 29.7%, respectively), consistent with active proliferation. Treatment with CPT-**2** reduced the EdU^+^ S phase population, in a concentration dependent manner, to the following values: 10.1% at 10 µM, 2.4% at 25 µM, and 1.1% at 50 µM (Fig. 4A and B). The CPT-**2** induced depletion of S phase cells was accompanied by a significant accumulation of cells in G2/M (42.6% at 25 µM and 35.3% at 50 µM, relative to 8.1% in DMSO). The G1 population did not change significantly (Fig. 4B). Treatment with **2** alone was indistinguishable from DMSO across all phases. The cell cycle profile produced by CPT-**2** is similar to Nutlin-3, which reduced the S phase population to 6.1%. At the highest CPT-**2** dose (50 µM), there is a very low population of cells in the S phase. The depletion of S phase cells by CPT-**2**, but not by **2**, demonstrates that the anti-proliferative effect requires CPT conjugation. Furthermore, the effects induced by CPT-**2** are consistent with a p53 dependent mechanism of cell cycle arrest downstream of MDM2-p53 disruption.

**Figure 4.**
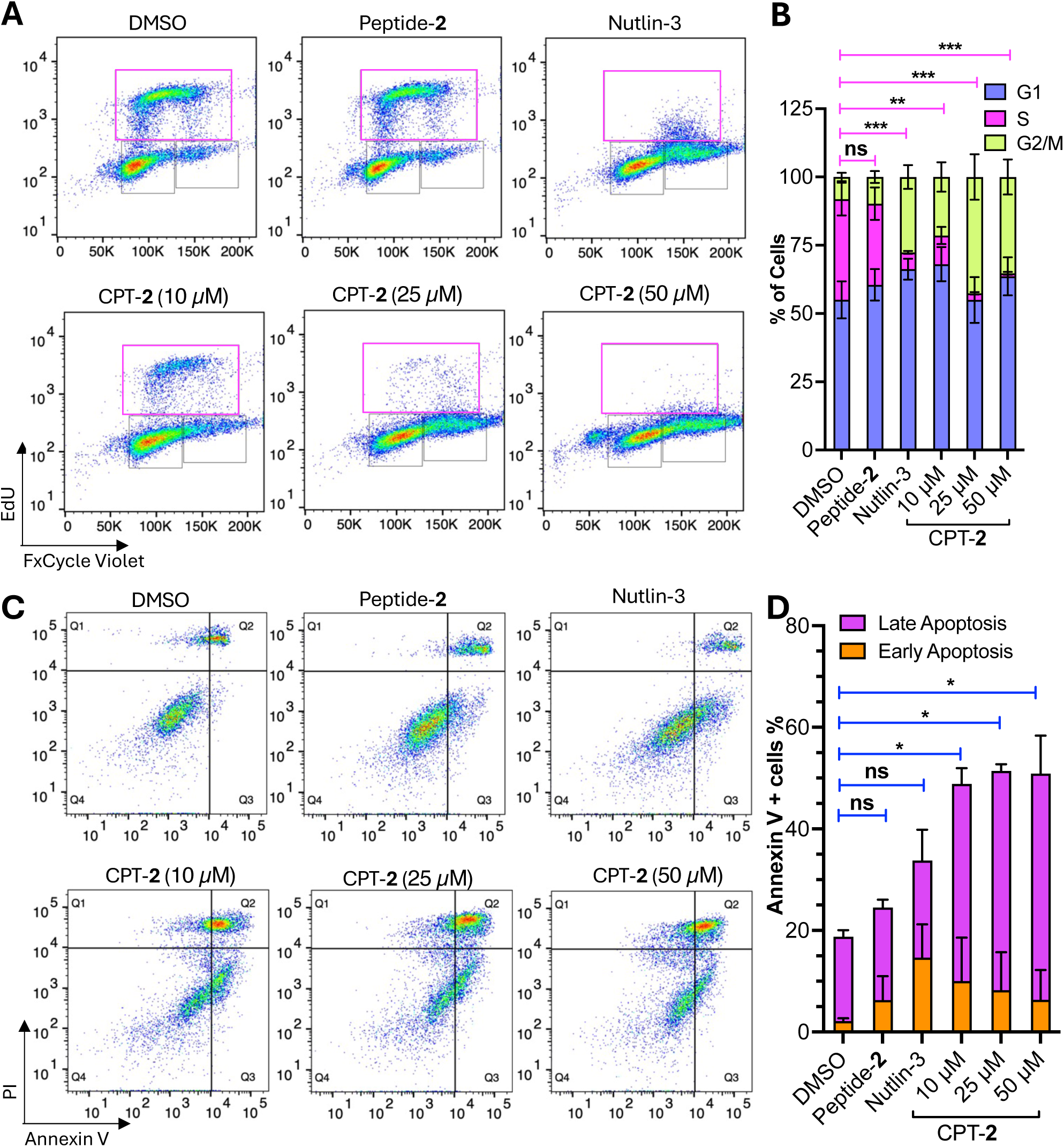
Effects of CPT-2 on cell cycle progression in MCF7 cells. **(A)** MCF7 cells were treated with DMSO (control), peptide-**2** (50 µM) alone, Nutlin-3 (10 µM), or increasing concentrations of CPT-**2** (10, 25, and 50 µM) for 24 h, followed by EdU incubation for an additional 2 h. Cell cycle distribution was analyzed by flow cytometry based on EdU incorporation (EdU-Alexa Fluor 488 staining) and DNA content (FxCycle Violet staining). Representative plots from three biological replicates are shown. **(B)** Quantification of the percentage of cells in each cell cycle phase following treatment. CPT-**2** induced an S-phase block in a concentration-dependent manner. Cell cycle distributions for the S phase were compared by two-way ANOVA followed by Dunnett’s multiple comparisons test versus vehicle (DMSO) control and are displayed above the bars; p ≤ 0.05 was considered significant. ***P≤ 0.001, **P= 0.0049, ns; not significant. **(C)** Representative flow cytometry plots of Annexin V-FITC and propidium iodide (PI) staining in MCF7 cells treated with DMSO, **2** (50 μM), Nutlin-3 (10 μM) or CPT-**2** (10, 25, or 50 μM) for 24 h. Cells were classified as viable (Q4, Annexin V^-^/PI^-^), early apoptotic (Q3, Annexin V^+^/PI^-^), late apoptotic (Q2, Annexin V^+^/PI^+^), or necrotic/dead (Q1, Annexin V^-^ /PI^+^). **(D)** Percentage of early (Q3, Annexin V^+^/PI^-^) and late apoptotic cell (Q2, Annexin V^+^/PI^+^) populations in response to the indicated treatments. Late apoptotic cell (Q2, Annexin V^+^/PI^+^) populations were compared by two-way ANOVA followed by Dunnett’s multiple comparisons test versus vehicle (DMSO) control and are displayed above the bars; p ≤ 0.05 was considered significant. *P is 0.0452 (DMSO vs 10 µM), 0.0160 (DMSO vs 25 µM) and 0.0118 (DMSO vs 50 µM). ns; not significant.

To determine whether the cells were also undergoing apoptosis, MCF7 cells were co-stained with Annexin V and propidium iodide (PI) after 24h of treatment and analyzed by flow cytometry (Fig. 4C and D). CPT-**2** increased the Annexin V positive population relative to DMSO. At CPT-**2** concentrations of 10, 25, and 50 µM, early apoptotic cells (Q3, Annexin V⁺/PI⁻) were quantified at 10%, 8.2%, and 6.4%, while late apoptotic cells (Q2, Annexin V⁺/PI⁺) were 38.8%, 43.2%, and 44.5%, respectively (Fig. 4C and D). In contrast, Nutlin-3 did not produce a significant increase in early or late apoptotic cells. Surprisingly, the total Annexin V positive fraction remained largely constant (∼49-51%) across the range of CPT-**2** concentrations; which contrasts with the results from cell-cycle arrest (Fig. 4B) and the loss of viability (Fig. 3C) which are dependent on CPT-**2** concentrations. It is possible that the Annexin V⁺/PI⁺ signal overestimates the true apoptotic fraction in this system. These findings indicate that engagement of the MDM2-p53 axis by CPT-**2** drives a predominantly cytostatic response that is concentration-dependent, and that cell death is more apparent at concentrations of 50 µM (Fig. 4C and D).

### Characterization of CPT cell uptake pathways

The chemical structure of CPT is very different from any other delivery vehicle that promotes cellular uptake. We therefore decided to investigate the pathways by which CPT-modified peptides enter cells. In the optimized CPT, the amines of the piperidine sidechains are likely protonated and positively charged under the aqueous conditions used for cell experiments. This series of positively charged secondary amines bears some resemblance to the frequent presence of lysine amino acids in cell penetrating peptides. The series of THF rings, however, are a very unexpected chemical feature that has not previously been used in delivery vehicles. To assess the contribution of THF rings to cellular uptake, a CPT analog lacking the THF groups (**S11**, Supporting Info, Table S3) was synthesized. This analog exhibited markedly reduced cellular uptake, with only ∼17% of MCF7 cells showing uptake of **S10**, thereby demonstrating that the THF rings are critical for CPT activity (Fig. S6A). Interestingly, unconjugated Fl-CPT (**S12**, Supporting Info, Table S3) was also efficiently internalized by MCF7 cells, exhibiting uptake levels comparable to those of Fl-CPT-**2** (∼95%). However, the MFI of Fl-CPT was significantly lower than that of Fl-CPT-**2** (Fig. S6B and C), indicating reduced intracellular accumulation despite similar uptake efficiency.

Next, the cellular uptake of Fl-CPT-**2** was evaluated to determine whether internalization in MCF7 cells occurs through an active transport mechanism. MCF7 cells were pretreated with either a combination of sodium azide and 2-deoxyglucose or incubated at 4 °C for 30 min to inhibit energy-dependent uptake pathways, followed by incubation with Fl-CPT-**2** for 3 h. Both conditions reduced Fl-CPT-**2** uptake by more than 50%, indicating that cellular internalization requires an energy-dependent active transport mechanism (Fig. 5A and B).

**Figure 5.**
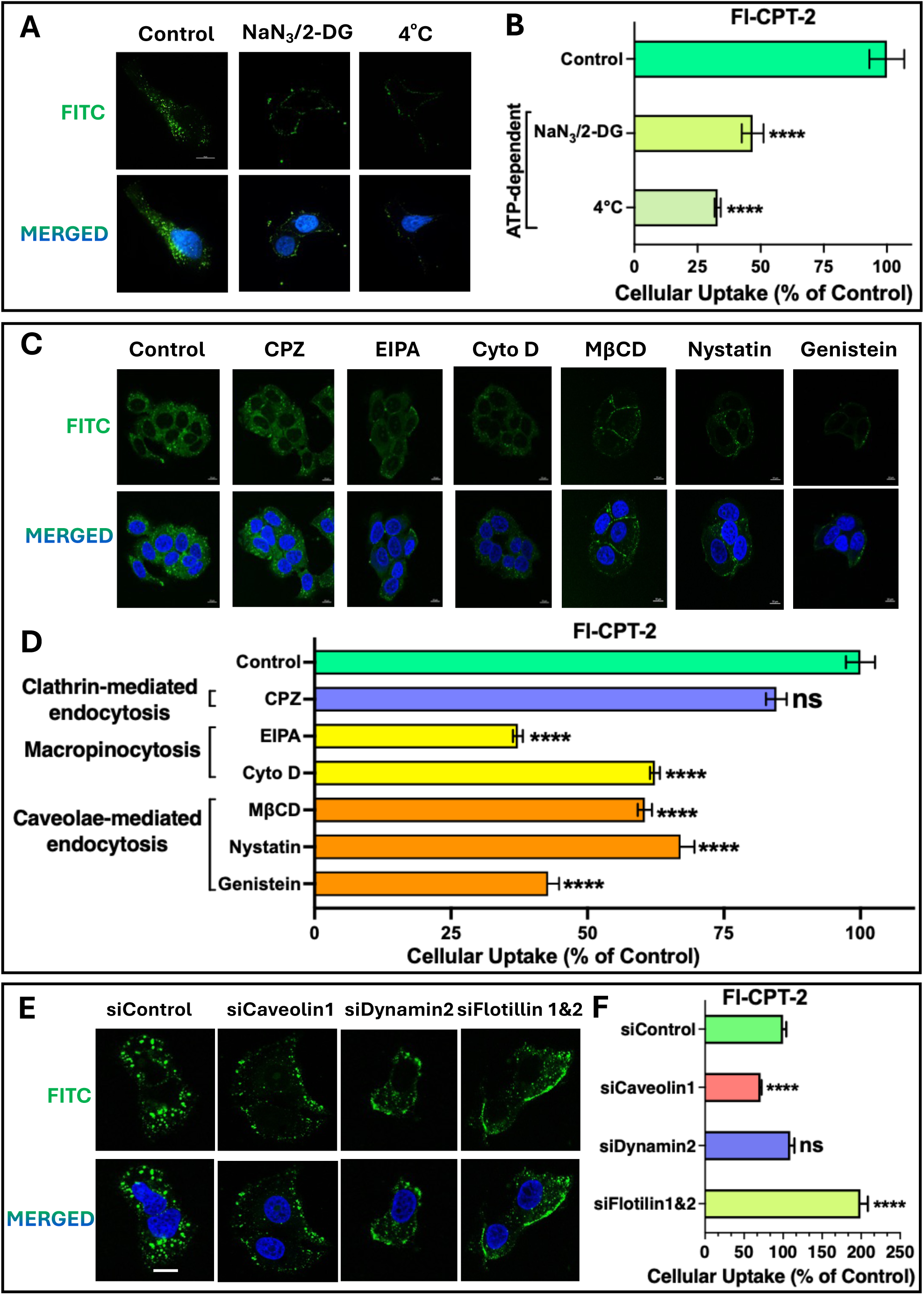
Mechanistic analysis of intracellular CPT-2 uptake in MCF7 cells. (**A**) Representative fluorescence microscopy images of MCF7 cells pretreated with NaN₃/2-DG or incubated at 4°C, followed by treatment with 5 µM Fl-CPT-**2**. (**B**) Quantification of intracellular Fl-CPT-**2** uptake under the indicated conditions, normalized to DMSO-treated control cells and presented as percentage uptake relative to control. Statistical significance was determined using the Kruskal-Wallis test, with p < 0.05 considered statistically significant. ****P ≤ 0.0001**. (C)** Representative fluorescence microscopy images of MCF7 cells pretreated with endocytic inhibitors prior to incubation with 5 µM Fl-CPT-**2**. **(D)** Quantification of intracellular Fl-CPT-**2** uptake following treatment with the indicated endocytic inhibitors, normalized to DMSO-treated control cells. Statistical significance was determined using the Kruskal-Wallis test, with p < 0.05 considered statistically significant. ****P ≤ 0.0001, ns; not significant. **(E)** Representative fluorescence microscopy images of MCF7 cells transfected with the indicated siRNAs followed by treatment with 5 µM Fl-CPT-**2**. **(F)** Quantification of intracellular Fl-CPT-**2** uptake in cells transfected with the indicated siRNAs, normalized to control siRNA-treated cells. Statistical significance was determined using the Mann Whitney U test, with p < 0.05 considered statistically significant. ****P ≤ 0.0001, ***p=0.0002, ns; not significant.

Cells internalize extracellular molecules through several active endocytic pathways, including clathrin-mediated, caveolae-mediated, and flotillin-mediated endocytosis, as well as macropinocytosis.^42^ These pathways rely on distinct cellular components and regulatory factors for efficient uptake. Pharmacological inhibitors targeting specific pathway-associated proteins or processes can therefore be used to selectively disrupt individual internalization mechanisms. To determine which pathways are important for uptake of Fl-CPT-**2**, MCF7 cells were pretreated with an inhibitor for 30 min, followed by incubation with 5 µM of Fl-CPT-**2** for 1 h in the presence of an inhibitor. Immunofluorescence analysis demonstrated that inhibition of clathrin-mediated endocytosis with chlorpromazine (CPZ) slightly decreased (∼15.4%) the cellular uptake of Fl-CPT-**2** (Fig. 5C and D), indicating a minimal contribution of this pathway to cellular internalization. In contrast, inhibition of macropinocytosis significantly decreased Fl-CPT-**2** uptake in MCF7 cells, with cytochalasin D and EIPA reducing uptake by >30% and ∼60%, respectively. These findings suggest that macropinocytosis represents one of the major routes of Fl-CPT-**2** entry into cells, consistent with the enhanced uptake capacity of tumor cells for extracellular material.^43^ Inhibition of caveolae-mediated endocytosis markedly impaired Fl-CPT-**2** internalization as well. Treatment with methyl-β-cyclodextrin (MβCD), genistein, and nystatin reduced cellular uptake by approximately 39.5%, 33%, and 57%, respectively, supporting the involvement of lipid raft associated endocytic pathways in Fl-CPT-**2** uptake (Fig. 5C and D).

Because MβCD and nystatin disrupt cholesterol-rich lipid rafts that are involved in multiple endocytic pathways, additional validation experiments were performed using transient knockdown of caveolin-1, dynamin-2, and flotillin-1&2 in MCF7 cells for 72 h and followed by incubation with Fl-CPT-**2** (Fig. S7A). After knockdown of caveolin-1 (a key structural component required for caveolae formation), Fl-CPT-**2** uptake was reduced by approximately 29%, further supporting the contribution of caveolae-mediated internalization. In contrast, knockdown of dynamin-2 produced only a minimal decrease in uptake (∼7.3%). Notably, silencing of flotillin-1 and flotillin-2 increased Fl-CPT-2 uptake by approximately 53%, suggesting that flotillin-dependent pathways are unlikely to assist with Fl-CPT-**2** internalization and may instead negatively regulate its uptake (Fig.5E and F).^44^ These observations were all consistent with imaging flow cytometry analysis (fig. S7B and C).

## Discussion

CPTs are a unique and versatile chemical scaffold to promote the delivery of peptide cargo into cells. A benefit of the CPT platform is the ability to easily modify sidechains with different chemical functionalities that alter cellular uptake properties (Figure S1). After exploring the uptake of CPTs with 8 different sidechains, an optimized CPT was developed for the delivery of either a polar or a hydrophobic peptide. It is surprising that the chemical combination of THF and piperidine sidechains leads to a very efficient CPT for cellular uptake as neither chemical group has previously been associated with cell penetrating peptides. These molecular features represent a significant departure from long-established observations of the chemical properties that lead to cellular uptake of cell penetrating peptides and other oligomeric synthetic scaffolds. Guanidine and amine groups (usually presented from arginine or lysine amino acids) are usually considered critical chemical entities for cell uptake, and many cell penetrating peptides as well as other related types of synthetic scaffolds rely on the presence of multiple guanidine and amine groups to promote cell uptake.^45,46^ Yet, endosomal entrapment frequently limits the bioactivity of cargo molecules conjugated to polycationic delivery agents.^6^ After CPT conjugation to peptides **1** and **2**, significantly enhanced cell uptake without endosomal entrapment was clearly demonstrated. CPT-**1** entered cells, showed diffuse distribution throughout the cytoplasm, but did not demonstrate reliable bioactivity, possible due to instability of the peptide or weak inhibitory activity. Extensive studies with CPT-**2** show that cell uptake and biological activity is greatly enhanced compared to peptide **2** alone, demonstrating that CPT-**2** can both enter cells and sufficiently escape from endosomes to exert biological activity. Investigating the uptake pathway of CPT-**2** showed that energy-dependent endocytosis is necessary, and that a combination of both macropinocytosis and caveolar-mediated endocytosis are dominant pathways. Interestingly, clathrin-mediated endocytosis is not important for uptake.

Peptide **2** had been previously optimized to inhibit MDM2 and MDMX,^29^ two proteins often overexpressed in cancer cells which inhibit the activity of p53. Many studies have focused on disrupting MDM2 binding to p53, and several clinical candidates have been examined.^47^ Despite the very potent inhibitory activity of **2** in vitro, the lack of cellular uptake and bioavailability likely hindered the further development of this peptide. This situation is common with many bioactive peptides as well as larger molecules that are unable to enter cells on their own. By coupling an optimized CPT with **2**, cellular delivery and biological activity is greatly enhanced. Relative to a small molecule positive control (Nutlin-3), CPT-**2** is also less toxic. These beneficial properties could reignite further clinical development of **2** as its CPT conjugate. The strategy of using CPT to enhance cellular uptake and biological activity can be applied to other molecules, making this an exciting new approach to advance the therapeutic development of peptides and other classes of molecules that have low bioavailability.

## Acknowledgments

We thank John Lloyd (NIDDK/ NIH) for performing mass spectrometry and David Taylor for assistance with NMR spectrometry.

## Author contributions

D.H.A., H.Z. and G.A. conceived and designed the experiments. H.Z. and H.A. prepared and purified all thyclotide monomers. H.Z. prepared and purified all peptides, thyclotides, and peptide-thyclotide conjugates. G.A. performed all cell experiments. V.C. performed a preliminary exploration of cell uptake experiments. A.T. conducted the data analysis of confocal microscopy images. F.L. conducted and analyzed Amnis imaging flow cytometry experiments. G.A. and M.K. performed the super-resolution microscopy experiments. D.H.A., H.Z., and G.A. cowrote the paper. D.H.A. supervised the entire study. All authors contributed to the final version of the paper.

## Disclaimer

This research was supported (in part) by the Intramural Research Program of the National Institute of Diabetes and Digestive and Kidney Diseases (NIDDK) within the National Institutes of Health (NIH). The contributions of the NIH author(s) are considered Works of the United States Government. The findings and conclusions presented in this paper are those of the author(s) and do not necessarily reflect the views of the NIH or the U.S. Department of Health and Human Services

## Supplementary Materials

### Chemicals and antibodies

Chlorpromazine (CPZ; Sigma-Aldrich, Cat. C8138) was dissolve in sterile water at a stock concentration of 30 mg/ml. Cytochalasin D (Sigma-Aldrich, Cat. C8273) was dissolved in DMSO to prepare a 5 mM stock solution and used at a final concentration of 2 µM. EIPA (5-(N-ethyl-N-isopropyl) amiloride; Selleckchem, Cat. S9849, 10 mM in DMSO) used at a final concentration of 100 µM. Methyl-ß-cyclodextrin (MßCD; Sigma-Aldrich, Cat. C4555) was dissolved in sterile cell culture grade water and applied at a final concentration of 10mM. Genistein (Sigma-Aldrich, Cat. G6649) was dissolved in DMSO and used at a final concentration of 200 µM. Nutlin-3 (Selleckchem, Cat. S1061) was dissolved in DMSO and used at a final concentration of 10 µM as a positive control for MDM2 inhibition.

For siRNA transfection, we used caveolin-1 siRNA(h) (Santa Cruz, Cat. sc-29241), Dynamin II siRNA(h) (Santa Cruz, Cat. sc-35236), Flotillin-1 siRNA(h) (Santa Cruz, Cat. sc-35391), Flotillin-2 siRNA(h) (Santa Cruz, Cat. sc-35393) and Control siRNA (Invitrogen, #AM4611).

For western blot, we used p53 (Calbiochem, Cat. OP43L; 1:1000), MDM2 (Santa Cruz, Cat. sc-965; 1:200), MDMX (Proteintech #84534-6-RR), p21 (Santa Cruz, Cat. sc-187; 1:200), BAX (Cell Signaling, Cat. 5023S; 1:1000), GAPDH (Cell Signaling Technology, Cat. D4C6R; 1:1000), caveolin-1 (Cell Signaling Technology, Cat. 3267S; 1:1000), Dynamin II (Santa Cruz, Cat. sc-166669; 1:200), Flotillin-1 (Santa Cruz, Cat. sc-74566; 1:200), Flotillin-2 (Santa Cruz, Cat. sc-28320; 1:200), γ-H2AX (Cell Signaling Technology, Cat. 5438S; 1:1000). For PLA experiment, we used p53 (Calbiochem, Cat. OP43L; 1:100) and MDM2 (Cell Signaling Technology, Cat. 86934; 1:100).

### Cell lines and cell culture

MCF7, MDA-MB-231 and SK-N-AS cells were purchased from American Type Culture Collection (ATCC, Manassas, VA, USA). All cell lines were maintained at 37 °C in a humidified incubator with 5% CO₂. MCF7 cells were cultured in EMEM supplemented with 10% FBS and 1% penicillin/streptomycin. SK-N-AS cells were cultured in DMEM supplemented with 10% FBS, 0.1 mM non-essential amino acids (NEAA), and 1% penicillin/streptomycin, while MDA-MB-231 cells were cultured in DMEM supplemented with 10% FBS and 1% penicillin/streptomycin. For treatments, all molecules were diluted in the respective complete growth medium for each cell line.

### Imaging Flow Cytometry

Cells were seeded in 6-well plates and, after 24 h, treated with fluorescein-labeled CPT-peptides diluted in culture medium. 16 hours after treatment, the peptide-containing medium was removed, and cells were washed three times with ice-cold PBS to prevent further uptake. Cells were detached using trypsin-EDTA and centrifuged at 1500 rpm for 5 min. Pellets were resuspended in 50 µl PBS containing 2% FBS and 20 µM DRAQ5, a DNA-staining dye (Abcam, Cat. ab108410). Cell internalization was measured on a 2-camera/12-channel Amnis ImageStream Mark II imaging flow cytometer (Cytek Bioscience, Fremont, CA) at 40X magnification equipped with 4 lasers. Data were recorded in Channel 1 (bright field), channel 2 (FITC) and channel 11 (DRAQ5) for 10,000 single cells per sample. For each cell, bright field (Channel 1), FITC (Channel 2) and DRAQ5 (Channel 11) images were used for the analysis. IDEAS 6.3 software (Amnis Corp, Seattle, WA) was used to analyze cellular uptake and intracellular distribution.

### Confocal Microscopy

MCF7 cells were seeded in µ-Slide 8-well chambers (Ibidi, Cat. 80806) at a density of 2 x10⁴ cells per well in complete medium. After 24 h, cells were treated with 5 µM fluorescein-labeled CPT-peptides diluted in complete medium. Following incubation 1, 3 or 16 h, depending on the experiment, cells were washed three times with ice-cold PBS and stained using Biotium Membrite Fix 640/660 Cell Surface Staining Kit according to the manufacturer’s instructions prior to fixation. Cells were then fixed with 4% formaldehyde (Thermo Fisher Scientific, Cat. 28906) for 15 min at room temperature, washed again with PBS, and mounted using SlowFade™ Diamond Antifade Mountant with DAPI (Thermo Fisher Scientific, Cat. S36973). For early endosome marker colocalization, cells were treated with CellLight BacMam 2.0 early endosome-RFP marker, Rab5a (Invitrogen, Carlbad, CA, USA) diluted in the culture medium. The endosome marker transduction was performed at a PPC of 50. Super-resolution images were acquired using either a Nikon SoRa spinning disk confocal microscope (Nikon Instruments Inc., Melville, NY, USA) equipped with a 60x apochromat oil immersion objective (NA 1.49) and a Hamamatsu ORCA Fusion BT sCMOS camera (Teledyne Photometrics, Tucson, AZ, USA), or a Zeiss LSM 880 confocal microscope with a 63x plan-apochromat oil immersion objective lens (NA 1.4) and Airyscan detector (Carl Zeiss Microscopy GmbH, Jena, Germany). Image deconvolution was performed using Nikon Elements software (v5.3) with a modified Richardson-Lucy iterative algorithm, or Zeiss ZEN Blue software for Airyscan processing.

### Western blotting

MCF7 and MDA-MB-231 cells were seeded in 6-well plates and treated with 10 µM Peptide-**2**, CPT-**2** or Nutlin-3 for 24 h. Cells were then washed with PBS, trypsinized, and lysed in RIPA buffer (Thermo Fisher Scientific, Cat. 89901) supplemented with a protease inhibitor cocktail (Roche, Cat.11836153001). Lysates were sonicated for 5 min (30 sec on/30 sec off cycles) at 4°C, and protein concentrations were determined using the Bradford assay (Thermo Fisher Scientific, Cat. 23200). Equal amounts of protein (30 µg) were resolved on 4-20% Mini-PROTEAN TGX precast gels (Bio-Rad, Cat. 4561096) and transferred onto nitrocellulose membranes (Bio-Rad, Cat. 1704159). Membranes were blocked for 1 hour in 5% non-fat milk prepared in TBS with 0.1% Tween-20 (TBS-T), then incubated overnight at 4°C with primary antibodies diluted in the blocking buffer. After three washes with TBS-T, membranes were incubated with HRP-conjugated secondary antibodies for 1 hour at room temperature. Protein bands were visualized using SuperSignal™ West Femto Maximum Sensitivity Substrate (Thermo Fisher Scientific, Cat. 34096) with a ChemiDoc MP Imaging System (Bio-Rad) and analyzed using Image Lab software (v6.1).

### Cytotoxicity Assay

Cell viability was determined using the Promega CellTiter 96® AQueous One Solution Cell Proliferation Assay (MTS) following the manufacturer’s protocol. Cells were seeded in 96-well tissue culture plates and treated the next day with DMSO, peptide-**2**, CPT alone, or CPT-**2** at serial concentrations ranging from 0.1 to 100 µM. After 72 hours of incubation, MTS reagent was added directly to each well and the plates were incubated for an additional 3 hours at 37 °C with 5% CO₂. Absorbance was measured at 490 nm using an Omega 640 spectrophotometer. Cell viability was calculated relative to DMSO-treated controls and averaged from three replicates.

### Proximity ligation assay (PLA)

PLA was performed using Duolink® In Situ Detection Reagents Red (Sigma, Cat. DUO92008-100RXN) according to the manufacturer’s instructions. Briefly, 2.5 x10⁴ MCF7 cells were seeded on Millicell EZ slides (Sigma, Cat. PEZGS0816) and cultured for 24 h. Cells were fixed with 4% paraformaldehyde in PBS for 15 min, permeabilized with 0.3% Triton X-100 for 10 min, and blocked with Duolink blocking solution at 37 °C for 1 h. Samples were then incubated overnight at 4 °C with primary antibodies against p53 and MDM2. Ligation and amplification were performed according to the manufacturer’s protocol. The ligation step was performed for 30 min at 37 °C, followed by polymerization reaction for 100 min at 37 °C. The cells were mounted with mounting media with DAPI. Fluorescent signals were imaged using a Zeiss LSM 880 confocal microscope.

### Cell uptake assessment with Endocytosis Inhibitors

MCF7 cells were seeded in µ-Slide 8-well chambers (Ibidi, #80806) at a density of 2×10^4^ cells per well in complete growth medium and cultured for 24 hours. The following day, the medium was replaced with fresh complete medium containing 0.1% DMSO or specific endocytosis inhibitors, and the cells were preincubated for 30 minutes. Subsequently, fluorescence-labeled CPT-**2** was added to the cultures in the presence or absence of inhibitors. After 1 hour of incubation, the cells were washed three times with ice-cold PBS and processed for either confocal microscopy or flow cytometry analysis. For quantitative analysis, cells were segmented as nuclear and cytoplasm and quantified using a custom script written in Phyton (v.3.10), utilizing the Cellpose deep-learning AI package (PMID: 39939718). Custom segmentation models were trained using the NIH HPC Biowulf cluster (http://hpc.nih.gov)

### Cell Uptake assessment with siRNA Transfection

All siRNAs were purchased from Santa Cruz Biotechnology and are listed in section “Chemicals and Antibodies”. MCF7 cells were seeded in 6-well plates at a density of 1×10^5^ cells per well in complete growth medium and cultured for 24 hours. siRNA transfection was performed using Lipofectamine RNAiMAX (Invitrogen, Cat. #13778075) according to the manufacturer’s instructions, at a final siRNA concentration of 10 nM. After 72 hours of transfection, the cells were incubated with fluorescence-labeled CPT-**2** at 37 °C for 1 hour, washed three times with ice-cold PBS, and processed for confocal microscopy or flow cytometry analysis. Quantitative analysis was performed using the same experimental conditions as those applied for the cell uptake assay with endocytosis inhibitors.

### Statistical analysis

Statistical analyses were performed using GraphPad Prisim 11. Data are presented as mean ± SEM. Comparisons between two or more groups were performed with Mann-Whitney, Kruskal Wallis or two-way ANOVA followed by Dunnett’s multiple comparisons test. A p-value < 0.05 was considered statistically significant. The *, **, ***, and **** symbols indicate P≤0.05, P≤0.01, P≤0.001, and P≤0.0001, respectively.

## Supplementary Tables and Figures

**Table S1.**
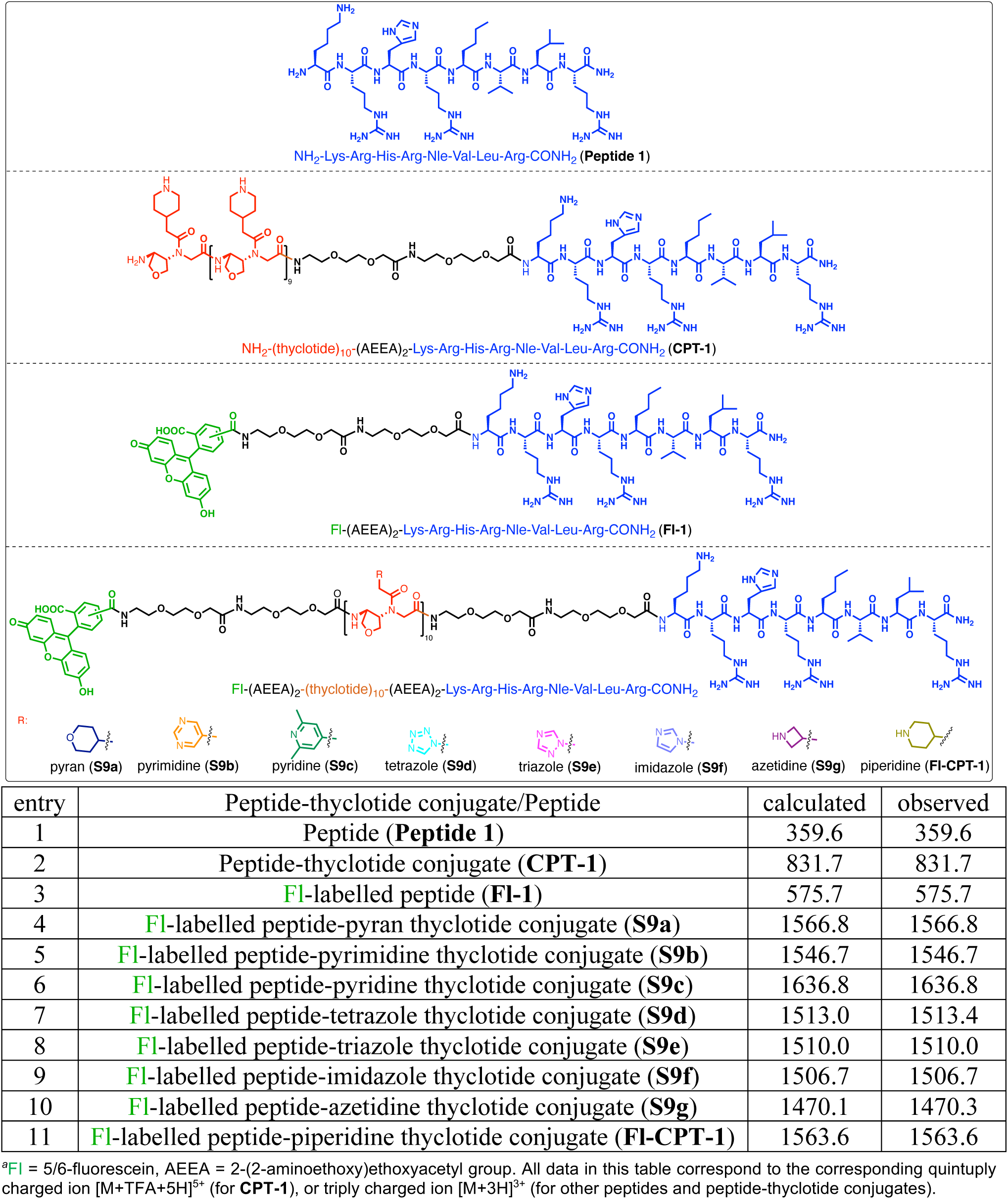
Mass characterization data for peptides and peptide-thyclotide conjugates (SETD8)*^a^*.

| NH<sub>2</sub>-Lys-Arg-His-Arg-Nle-Val-Leu-Arg-CONH<sub>2</sub> (**Peptide 1**) |  |  |  |
| --- | --- | --- | --- |
| NH<sub>2</sub>-(thyclotide)<sub>10</sub>-(AEEA)<sub>2</sub>-Lys-Arg-His-Arg-Nle-Val-Leu-Arg-CONH<sub>2</sub> (**CPT-1**) |  |  |  |
| FI-(AEEA)<sub>2</sub>-Lys-Arg-His-Arg-Nle-Val-Leu-Arg-CONH<sub>2</sub> (**FI-1**) |  |  |  |
| FI-(AEEA)<sub>2</sub>-(thyclotide)<sub>10</sub>-(AEEA)<sub>2</sub>-Lys-Arg-His-Arg-Nle-Val-Leu-Arg-CONH<sub>2</sub><br>R:<br>pyran (**S9a**)<br>pyrimidine (**S9b**)<br>pyridine (**S9c**)<br>tetrazole (**S9d**)<br>triazole (**S9e**)<br>imidazole (**S9f**)<br>azetidine (**S9g**)<br>piperidine (**FI-CPT-1**) |  |  |  |
| entry | Peptide-thyclotide conjugate/Peptide | calculated | observed |
| 1 | Peptide (**Peptide 1**) | 359.6 | 359.6 |
| 2 | Peptide-thyclotide conjugate (**CPT-1**) | 831.7 | 831.7 |
| 3 | FI-labelled peptide (**FI-1**) | 575.7 | 575.7 |
| 4 | FI-labelled peptide-pyran thyclotide conjugate (**S9a**) | 1566.8 | 1566.8 |
| 5 | FI-labelled peptide-pyrimidine thyclotide conjugate (**S9b**) | 1546.7 | 1546.7 |
| 6 | FI-labelled peptide-pyridine thyclotide conjugate (**S9c**) | 1636.8 | 1636.8 |
| 7 | FI-labelled peptide-tetrazole thyclotide conjugate (**S9d**) | 1513.0 | 1513.4 |
| 8 | FI-labelled peptide-triazole thyclotide conjugate (**S9e**) | 1510.0 | 1510.0 |
| 9 | FI-labelled peptide-imidazole thyclotide conjugate (**S9f**) | 1506.7 | 1506.7 |
| 10 | FI-labelled peptide-azetidine thyclotide conjugate (**S9g**) | 1470.1 | 1470.3 |
| 11 | FI-labelled peptide-piperidine thyclotide conjugate (**FI-CPT-1**) | 1563.6 | 1563.6 |

<sup>a</sup>FI = 5/6-fluorescein, AEEA = 2-(2-aminoethoxy)ethoxyacetyl group. All data in this table correspond to the corresponding quintuply charged ion [M+TFA+5H]<sup>5+</sup> (for **CPT-1**), or triply charged ion [M+3H]<sup>3+</sup> (for other peptides and peptide-thyclotide conjugates).

**Table S2.**
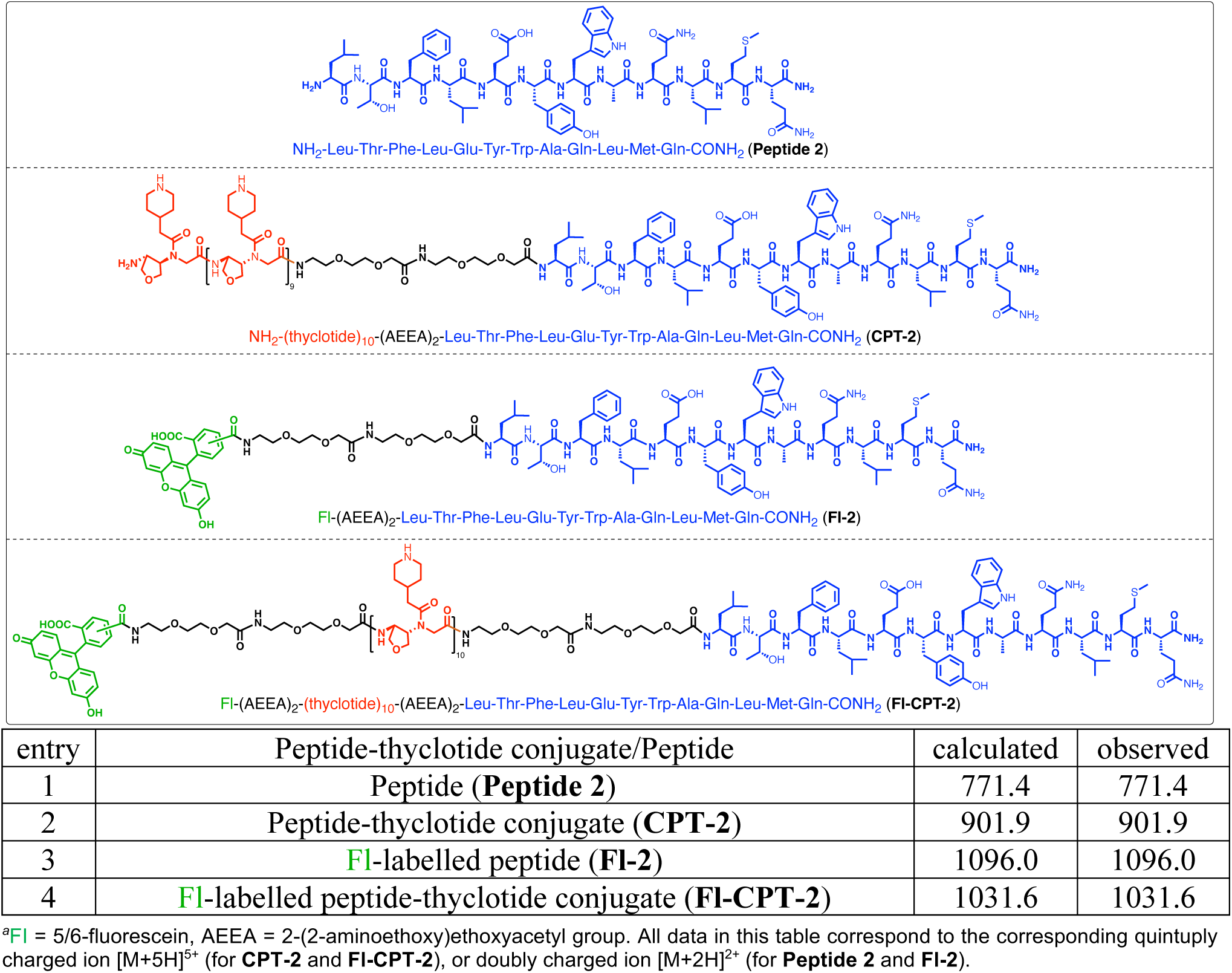
Mass characterization data for peptides and peptide-thyclotide conjugates (MDM2)*^a^*.

| NH<sub>2</sub>-Leu-Thr-Phe-Leu-Glu-Tyr-Trp-Ala-Gln-Leu-Met-Gln-CONH<sub>2</sub> (**Peptide 2**) |  |  |  |
| --- | --- | --- | --- |
| NH<sub>2</sub>-(thyclotide)<sub>10</sub>-(AEEA)<sub>2</sub>-Leu-Thr-Phe-Leu-Glu-Tyr-Trp-Ala-Gln-Leu-Met-Gln-CONH<sub>2</sub> (**CPT-2**) |  |  |  |
| FI-(AEEA)<sub>2</sub>-Leu-Thr-Phe-Leu-Glu-Tyr-Trp-Ala-Gln-Leu-Met-Gln-CONH<sub>2</sub> (**FI-2**) |  |  |  |
| FI-(AEEA)<sub>2</sub>-(thyclotide)<sub>10</sub>-(AEEA)<sub>2</sub>-Leu-Thr-Phe-Leu-Glu-Tyr-Trp-Ala-Gln-Leu-Met-Gln-CONH<sub>2</sub> (**FI-CPT-2**) |  |  |  |
| entry | Peptide-thyclotide conjugate/Peptide | calculated | observed |
| 1 | Peptide (**Peptide 2**) | 771.4 | 771.4 |
| 2 | Peptide-thyclotide conjugate (**CPT-2**) | 901.9 | 901.9 |
| 3 | FI-labelled peptide (**FI-2**) | 1096.0 | 1096.0 |
| 4 | FI-labelled peptide-thyclotide conjugate (**FI-CPT-2**) | 1031.6 | 1031.6 |

<sup>a</sup>FI = 5/6-fluorescein, AEEA = 2-(2-aminoethoxy)ethoxyacetyl group. All data in this table correspond to the corresponding quintuply charged ion [M+5H]<sup>5+</sup> (for **CPT-2** and **FI-CPT-2**), or doubly charged ion [M+2H]<sup>2+</sup> (for **Peptide 2** and **FI-2**).

**Table S3.**
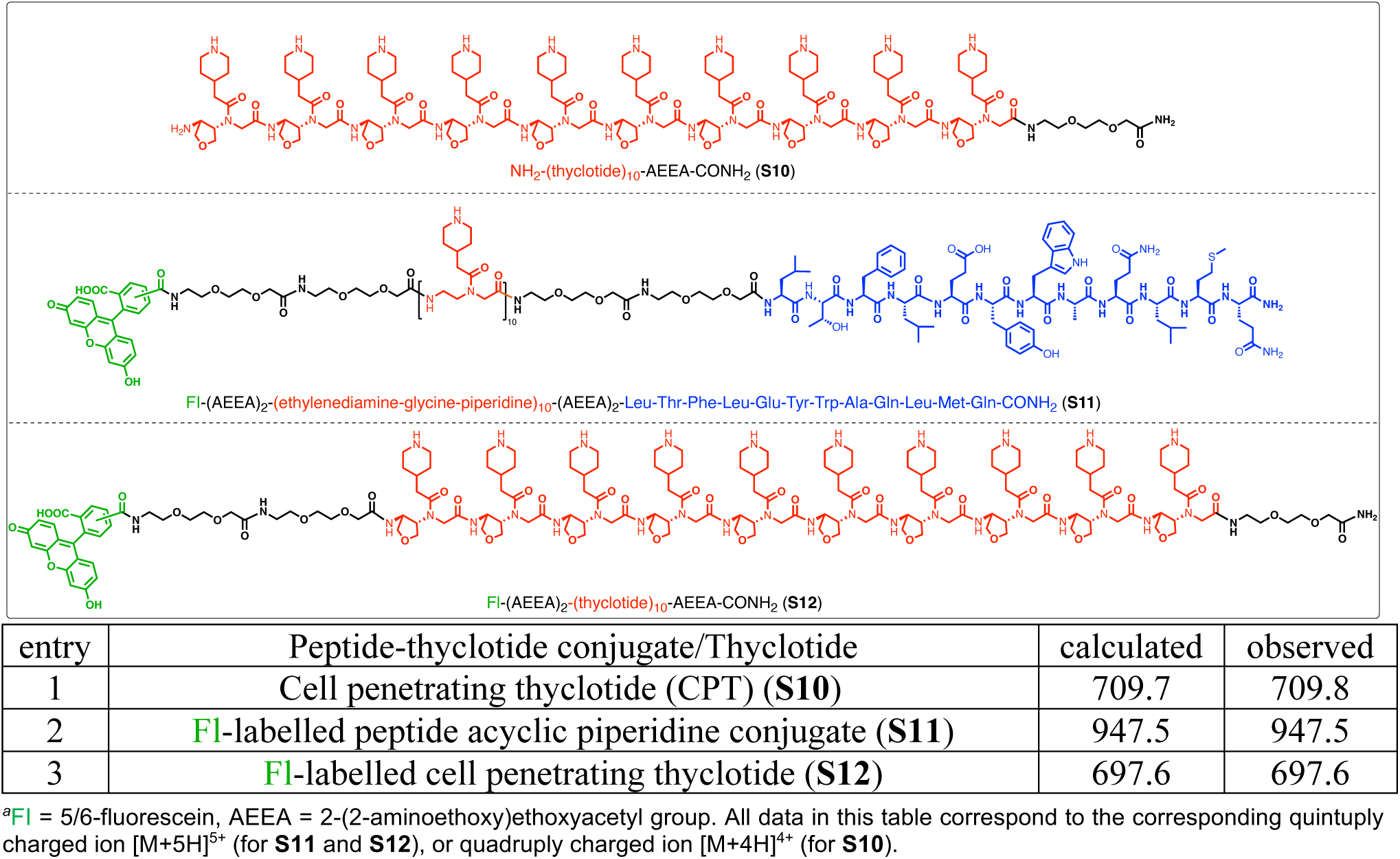
Mass characterization data for negative control molecules*^a^*.

| NH<sub>2</sub>-(thyclotide)<sub>10</sub>-AEEA-CONH<sub>2</sub> (**S10**) |  |  |  |
| --- | --- | --- | --- |
| FI-(AEEA)<sub>2</sub>-(ethylenediamine-glycine-piperidine)<sub>10</sub>-(AEEA)<sub>2</sub>-Leu-Thr-Phe-Leu-Glu-Tyr-Trp-Ala-Gln-Leu-Met-Gln-CONH<sub>2</sub> (**S11**) |  |  |  |
| FI-(AEEA)<sub>2</sub>-(thyclotide)<sub>10</sub>-AEEA-CONH<sub>2</sub> (**S12**) |  |  |  |
| entry | Peptide-thyclotide conjugate/Thyclotide | calculated | observed |
| 1 | Cell penetrating thyclotide (CPT) (**S10**) | 709.7 | 709.8 |
| 2 | FI-labelled peptide acyclic piperidine conjugate (**S11**) | 947.5 | 947.5 |
| 3 | FI-labelled cell penetrating thyclotide (**S12**) | 697.6 | 697.6 |

<sup>a</sup>FI = 5/6-fluorescein, AEEA = 2-(2-aminoethoxy)ethoxyacetyl group. All data in this table correspond to the corresponding quintuply charged ion [M+5H]<sup>5+</sup> (for **S11** and **S12**), or quadruply charged ion [M+4H]<sup>4+</sup> (for **S10**).

**Fig. S1.**
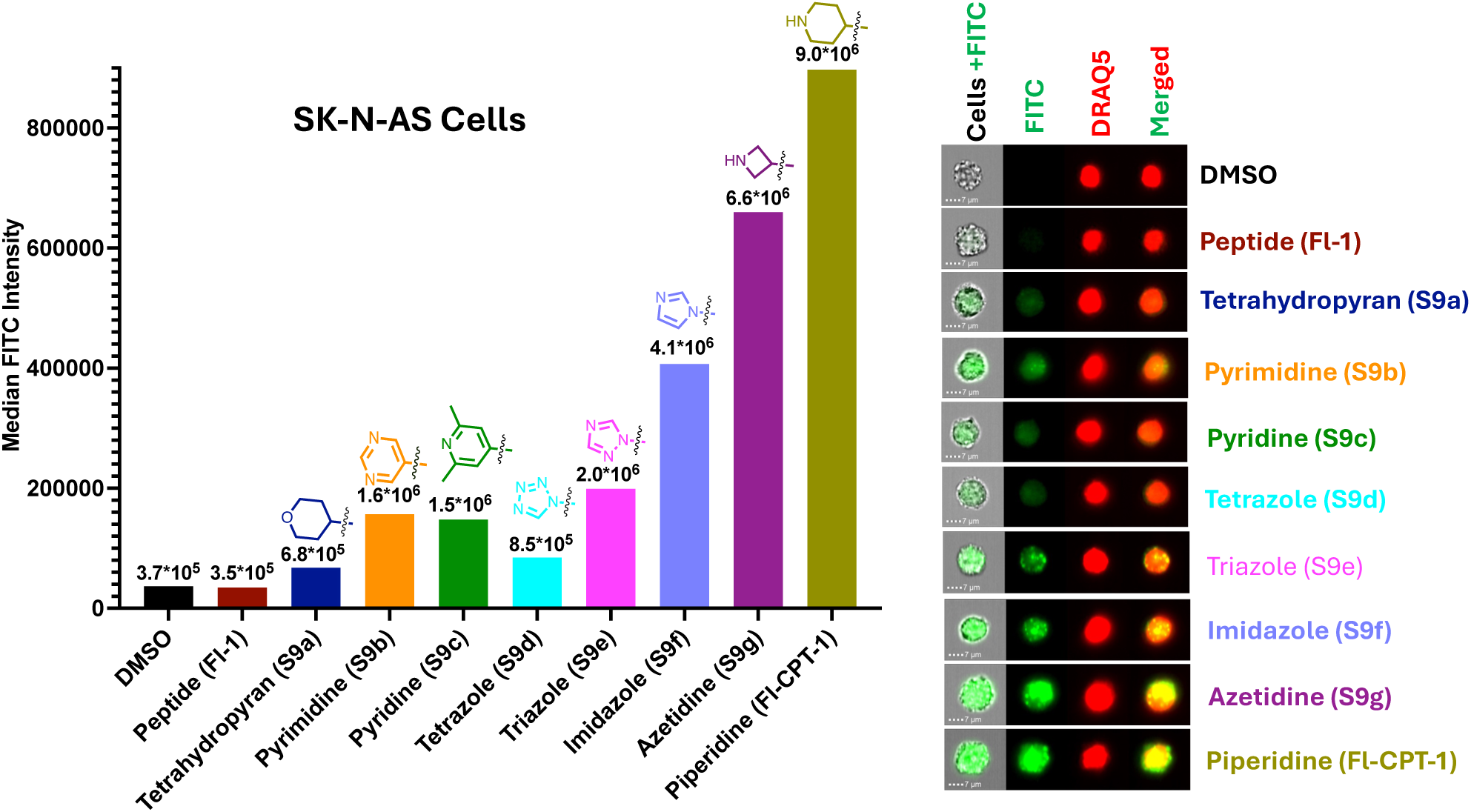
Screening of CPT conjugates containing different functional moieties. **(A)** SK-N-AS cells were treated with 5 µM peptide-**1** conjugated to different CPT derived moieties and analyzed by imaging flow cytometry to evaluate intracellular uptake based on median fluorescence intensity (MFI). **(B)** Representative imaging flow cytometry images of cells treated with each conjugate. Nuclei were stained with DRAQ5. Scale bar, 7 µm.

**Fig. S2.**
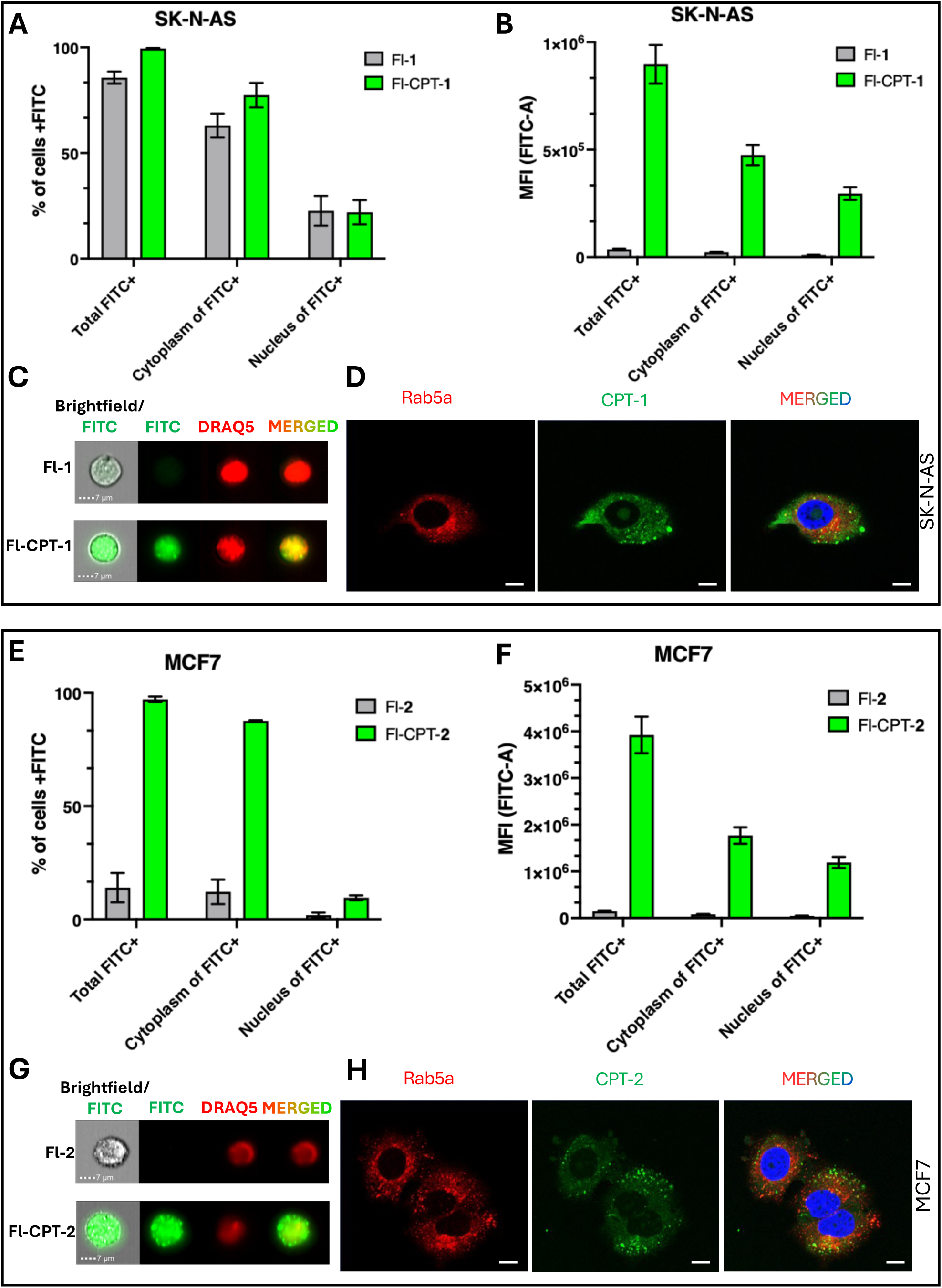
Cellular uptake efficiency of CPT-peptide conjugates. **(A)** Percentage of SK-N-AS cells positive for internalized Fl-**1** or Fl-CPT-**1** following treatment with 5 µM compounds for 16 h, as determined by imaging flow cytometry. **(B)** Quantification of intracellular uptake based on median fluorescence intensity (MFI). **(C)** Representative imaging flow cytometry images of SK-N-AS cells treated with Fl-**1** or Fl-CPT-**1**. Nuclei were stained with DRAQ5. **(D)** SK-N-AS cells were transfected with Rab5a (CellLight Early Endosomes-RFP) 24 h prior to incubation with Fl-CPT-**1** for 1 h. Nuclei were counterstained with DAPI. Scale bar, 10 µm. **(E)** Percentage of MCF7 cells positive for internalized Fl-**2** or Fl-CPT-**2** following treatment with 5 µM compounds for 16 h, as determined by imaging flow cytometry. **(F)** Quantification of intracellular uptake based on median fluorescence intensity (MFI). **(G)** Representative imaging flow cytometry images of cells treated with Fl-**2** or Fl-CPT-**2**. Nuclei were stained with DRAQ5. **(H)** MCF7 cells were transfected with Rab5a (CellLight Early Endosomes-RFP) 24 h prior to incubation with Fl-CPT-**2** for 1 h. Nuclei were counterstained with DAPI. Scale bar, 10 µm.

**Fig. S3.**
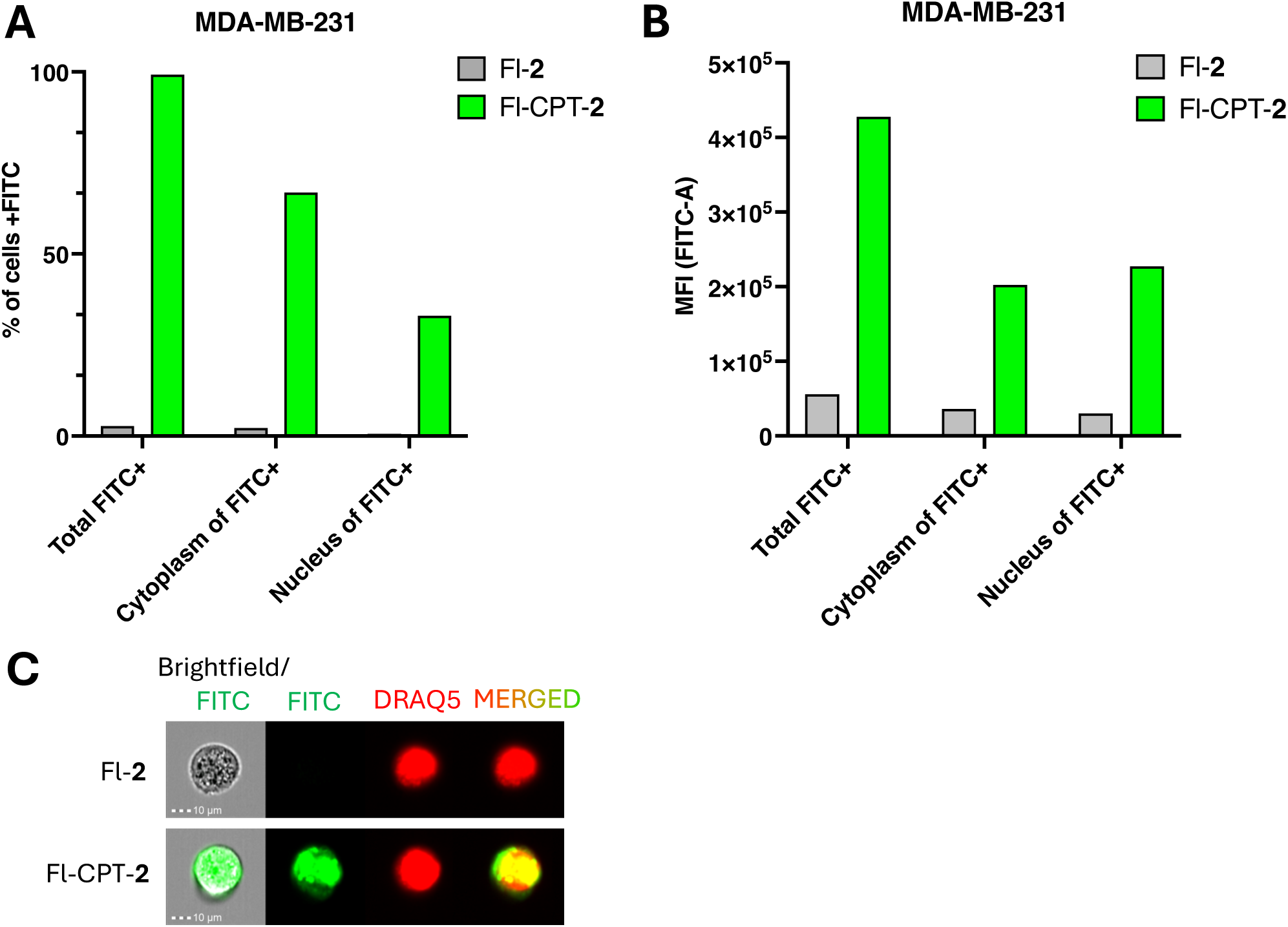
Cellular uptake efficiency of CPT-peptide-2 on MDA-MB-231 cells. **(A)** Percentage of MDA-MB-231 cells positive for internalized Fl-**2** or Fl-CPT-**2** following treatment with 5 µM compounds for 16 h, as determined by Amnis imaging flow cytometry. **(B)** Quantification of intracellular uptake based on median fluorescence intensity (MFI). **(C)** Representative imaging flow cytometry images of cells treated with Fl-**2** or Fl-CPT-**2**. Nuclei were stained with DRAQ5.

**Fig. S4.**
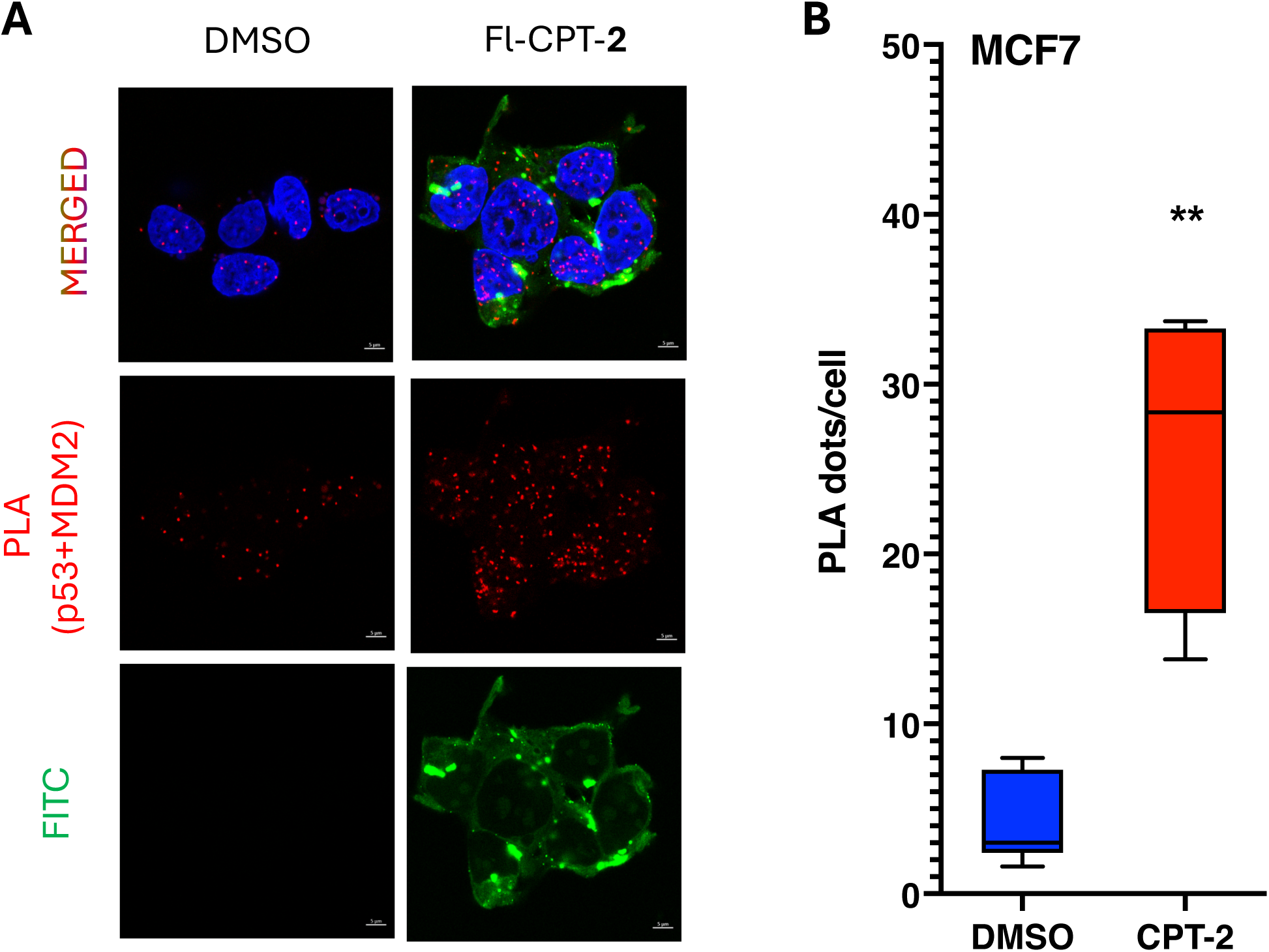
CPT-2 enhances nuclear p53-MDM2 interaction in MCF7 cells. **(A)** Proximity ligation assay (PLA) of MCF7 cells treated with DMSO or 10 µM CPT-**2** for 24 h. Red fluorescent puncta indicate close proximity and interaction between p53 and MDM2 within the nucleus. **(B)** Quantification of p53-MDM2 interactions in cells treated with DMSO or CPT-**2**. Data are presented as mean ± SD from 10 analyzed images from each sample. Statistical significance was determined using the Mann-Whitney test; **p < 0.01.

**Fig. S5.**
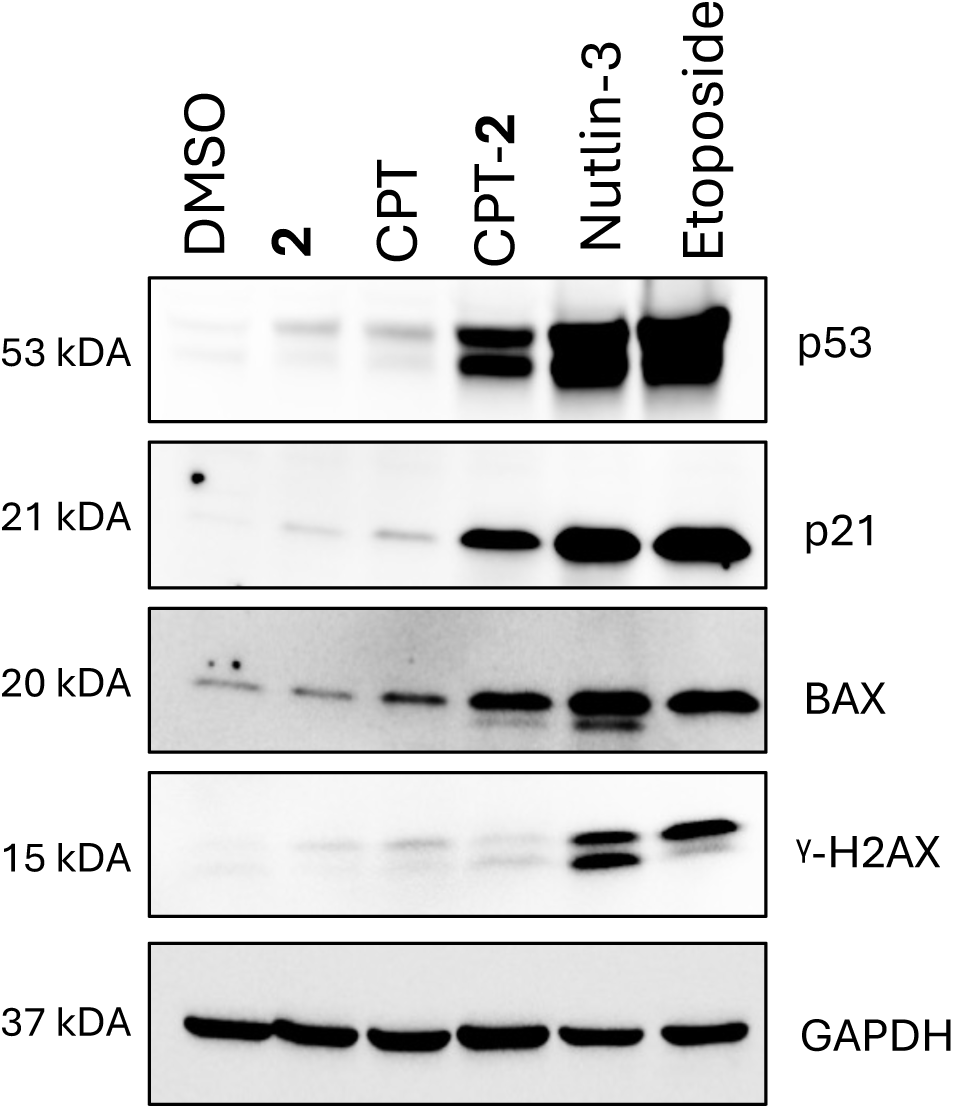
CPT-2 does not induce DNA damage response. Western blot analysis of MCF7 cells treated with the indicated compounds at 50 µM for 96 h, except for Nutlin-3, which was used at 10 µM. Representative western blots demonstrate changes in protein expression associated with prolonged treatment. Increased γ-H2AX staining indicates induction of DNA damage response pathways.

**Fig. S6.**
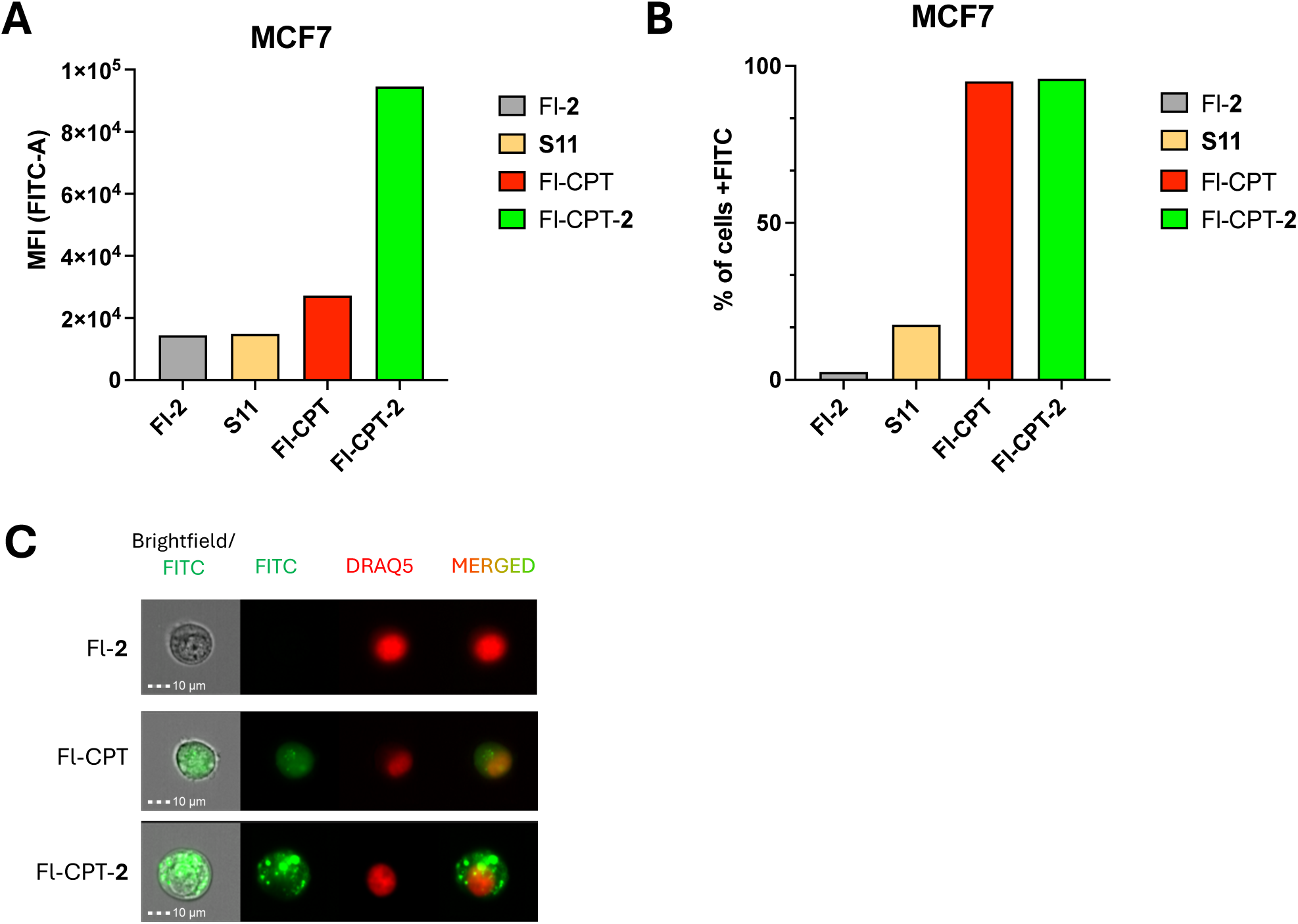
THF ring is critical for cellular uptake. **(A)** Percentage of MCF7 cells positive for internalized for Fl-**2**, **S11**, Fl-CPT and FL-CPT-**2** following treatment with 5 µM compounds for 16 h, as determined by Amnis imaging flow cytometry. **(B)** Quantification of intracellular uptake based on median fluorescence intensity (MFI). **(C)** Representative imaging flow cytometry images of cells treated with Fl-**2,** Fl-CPT (**S12**) or Fl-CPT-**2**. Nuclei were stained with DRAQ5.

**Fig. S7.**
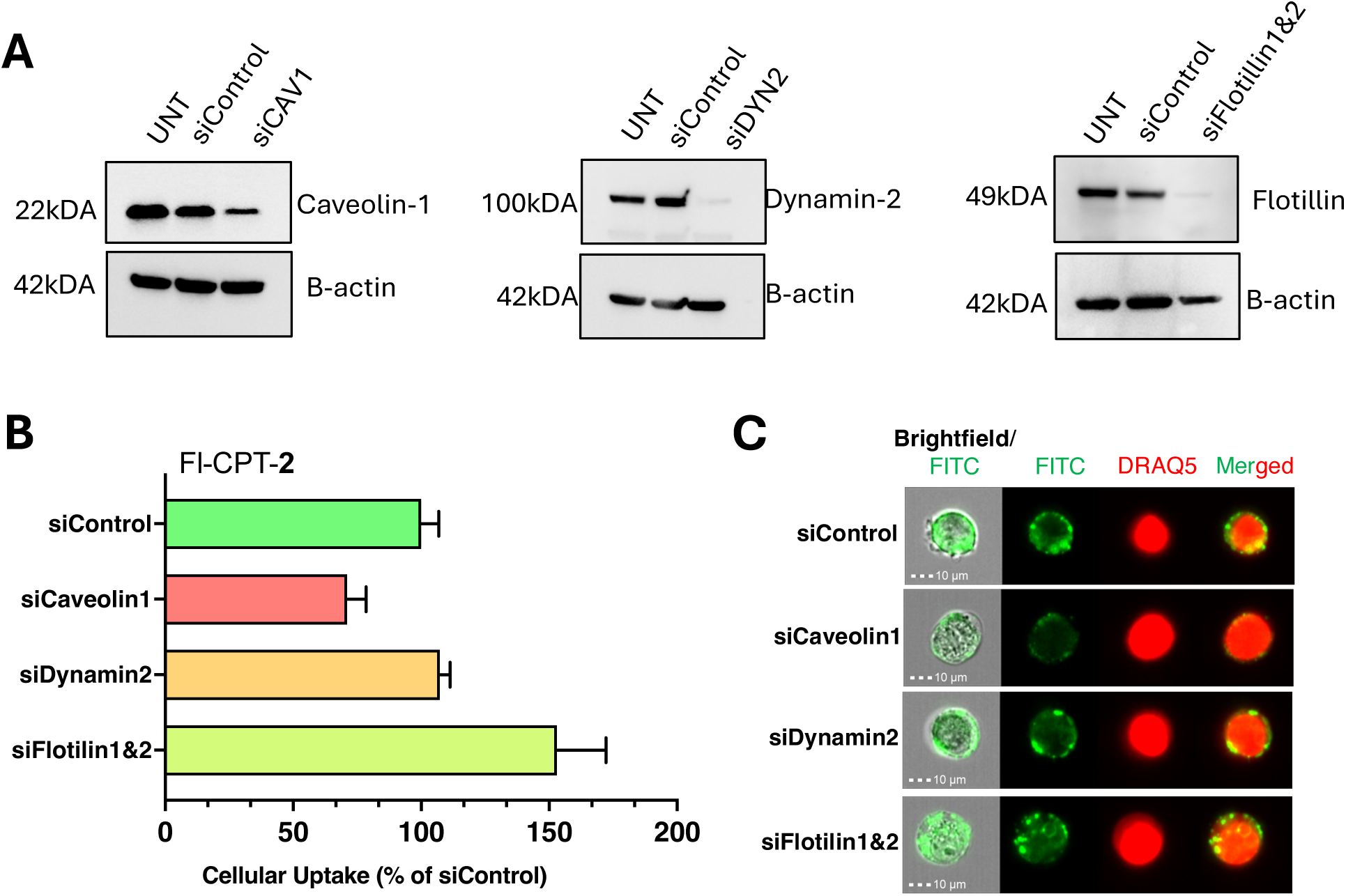
Assessment of lipid raft-mediated endocytosis following siRNA-mediated knockdown of target genes. **(A)** MCF7 cells were transiently transfected with 10 nM gene-specific siRNAs for 72 h. Knockdown efficiency was validated by western blot analysis using antibodies against Caveolin-1, Dynamin-2, Flotillin-1&2, and β-actin as a loading control. **(B)** Quantification of intracellular Fl-CPT-**2** uptake under the indicated knockdown conditions. Cellular uptake was analyzed by Amnis imaging flow cytometry, normalized to siControl-transfected cells, and expressed as the percentage relative to siControl. **(C)** Representative imaging flow cytometry images showing intracellular fluorescence intensity of Fl-CPT-**2** in MCF7 cells transfected with siControl, siCaveolin-1, siDynamin-2, or siFlotillin-1&2 for 72 h, followed by incubation with Fl-CPT-**2** for 1 h.

